# Integrated Transcriptomic Profiling Reveals MAPK14-Related Inflammatory Signalling Associated with Postoperative Atrial Fibrillation

**DOI:** 10.64898/2026.09.01.748407

**Authors:** Jiqiu Hou, Nicola Tidbury, Joshua Preston, Bilal H. Kirmani, Ethan Haynes, Nordine Helassa, Yuze Ma, Aiswarya Dev, Frances Greaney, Iain M. Dykes, Parveen Sharma, Gregory Y. H. Lip, Rebecca A. B. Burton

**Author notes:** Equal contributing first authors (Jiqiu Hou, Nicola Tidbury). Corresponding author – Dr Rebecca AB Burton.

## Abstract

Postoperative atrial fibrillation (POAF) affects up to half of patients after cardiac surgery and raises long-term risk of stroke and death. Why only some patients develop it, despite similar surgical exposure, remains poorly understood. We studied a pilot cohort of 20 patients undergoing cardiac surgery in Northwest England, comparing those who developed POAF within four days (n=10) with those who did not (n=10). Circulating plasma RNA was profiled using the NanoString nCounter Cardiovascular Disease panel, and differentially expressed genes were mapped onto protein-protein interaction networks using STRING and Cytoscape. Candidate genes were then validated by RT-qPCR in the same cohort, alongside serial ELISA measurement of plasma MAPK14 protein. Peripheral blood monocyte, neutrophil and total white blood cell counts were also compared between the two groups on postoperative Days 1, 2 and 4.

Screening identified MAPK14, COX6C and COX7B as elevated in POAF, with PTPN12 reduced. These genes clustered within a connected inflammatory and stress-response network, and pathway enrichment pointed to immune activation. RT-qPCR confirmed higher MAPK14, COX6C and COX7B expression in POAF patients. Plasma MAPK14 rose after surgery in both groups, with the largest increase in POAF patients on day two, though this did not reach significance. Peripheral blood monocyte counts were significantly higher in POAF patients than in those without POAF at all three postoperative timepoints, whereas neutrophil and total white blood cell counts did not differ significantly between groups.

These early findings identify a circulating plasma RNA signature linked to MAPK14-related inflammatory signalling as a candidate marker of POAF susceptibility, warranting confirmation in a larger, adequately powered cohort.

## Introduction

Postoperative atrial fibrillation (POAF) is one of the most common complications after cardiac surgery, occurring in an estimated 15–50% of patients and associated with an increased risk of stroke, heart failure and long-term mortality [1, 2]. POAF has traditionally been regarded as a transient response to surgical stress. Follow-up data from the VISION Cardiac Surgery study and the SWEDEHEART registry show that patients who develop POAF remain at substantially higher risk of AF recurrence for years afterwards [3, 4], which argues against a purely transient mechanism and points instead to an underlying atrial vulnerability that surgery exposes rather than creates. Among the proposed mechanisms, inflammation has been most consistently implicated in this vulnerability; specifically, macrophage-derived IL-6 signalling has been demonstrated to induce calcium leak in atrial myocytes [5], circulating markers such as C-reactive protein and neutrophil-to-lymphocyte ratio track with POAF risk [6]. Trials of colchicine, corticosteroids, posterior pericardiotomy and partial cardiac denervation have each shown that reducing perioperative inflammation or autonomic activation can lower POAF incidence, even though results have not been uniformly positive [7–10]. Circulating monocyte counts have themselves been reported to associate with POAF risk and early postoperative mortality after cardiac surgery [11], raising the possibility that monocyte-linked inflammatory cells contribute directly to this vulnerability.

What remains unclear is why only a subset of patients subjected to comparable surgical stress subsequently develop POAF. Prediction models built from ECG features, clinical variables and circulating biomarkers, including several using machine learning, can estimate POAF risk but are predictive rather than explanatory, and offer little insight into the underlying biology [12, 13]. To investigate this question, we studied a pilot cohort of patients undergoing cardiac surgery. We combined transcriptomic profiling using the NanoString nCounter platform with network and pathway analyses (STRING/Cytoscape) and RT-qPCR validation to determine whether POAF is associated with a distinct molecular signature. We further examined whether MAPK14-related inflammatory signalling, identified through the transcriptomic screen, could be validated as a marker of POAF susceptibility. We also examined whether this transcriptomic signal was accompanied by changes in circulating monocyte, neutrophil and white blood cell counts, as complementary cellular indices of the postoperative inflammatory response.

## Methods

### Ethics

This work formed part of the broader AF Big Picture project, Insights into the Underlying Pathophysiology of Atrial Fibrillation (REC 21/EE/0040; IRAS ID: 265408). The study adhered to the principles of the Declaration of Helsinki and followed established standards for ethical and inclusive research. All participant data were anonymised and stored securely in accordance with GDPR requirements. Written informed consent was obtained from all participants prior to tissue collection, and all human tissue was collected, stored, and handled in compliance with the Human Tissue Act 2004.

### Study population and POAF classification

Patients undergoing cardiac surgery (coronary artery bypass grafting and/or aortic/mitral valve repair or replacement) were recruited, and patients with no prior atrial fibrillation (AF) who developed POAF by day 4 (n=10) were compared with those who remained free of POAF (n=10) in the 4 days post-surgery; POAF was defined as any documented AF on days 1–4 post-surgery and this was based on ECG recordings. In the no POAF group, none of the patients developed new POAF between hospital discharge and six-to-eight-week follow-up. Baseline demographic, clinical and surgical characteristics were compared between the two groups. Continuous variables (age and BMI) were first assessed for normality using the Shapiro-Wilk test; both variables followed a normal distribution in each group and are presented as mean±SD. Between-group comparisons of continuous variables were performed using an unpaired t test with Welch’s correction, which does not assume equal variances between groups. Categorical variables (gender, smoking status, previous cardiovascular event, previous atrial fibrillation, beta-blocker usage, and type of surgery) are presented as counts and percentages, n (%), and were compared between groups using Fisher’s exact test, which is appropriate given the small sample size in each group. A two-tailed P value <0.05 was considered statistically significant. All statistical analyses were performed using GraphPad Prism (version 11.0.0, GraphPad Software, San Diego, CA, USA).

### Blood collection and plasma processing

4ml of EDTA plasma was collected immediately before surgery (Day 0) and days 1, 2 and 4 post-surgery in line with standard clinical care. Blood was centrifuged within 2 hours of collection and resultant plasma was stored at –80°C until analysis. Presence of POAF was determined using standard of care procedures and was documented in patient notes.

### RNA extraction and nCounter gene expression profiling

Total RNA was extracted from plasma using the Promega Maxwell RSC with the Maxwell® RSC miRNA from Tissue or Plasma and Serum Kit (Promega AS1680) and quantified on the nanodrop and qubit 3.0 fluorometer. A total amount of 100ng was hybridised with Reporter and Capture Probesets from the nCounter® Cardiovascular Disease (CVD) Pathophysiology panel and counted using the nCounter Pro. Samples passed quality control (QC) under the following parameters: field of view (FOV) registration >75%, Binding density between 0.1 and 2.25, and positive control R^2^ >0.95. Endogenous normalisers were identified within the dataset using the approach described by Adamova et al. [14]. Raw counts were processed in nSolver using six normalisation and background-correction workflows. Protein-protein interaction networks were constructed using STRING and visualised in Cytoscape, and functional enrichment analysis was performed to identify the biological processes represented among candidate genes, with particular attention to immune activation, inflammatory signalling, extracellular processes, stress response and mitochondrial function; enrichment bubble plots were generated in RStudio, R version 4.6.1.

### RT-qPCR validation of candidate genes

Candidate differentially expressed genes (DEGs), including MAPK14, COX6C, COX7B and PTPN12, were validated by RT-qPCR using UBB and PPIA as reference genes, in a validation cohort designed as n=6 per group (random selection from the cohort); primer sequences are provided in the Supplementary Table 1, and primers were purchased from Merck (Supplementary Table 1). Relative expression was calculated using the 2^−ΔΔCt^ method.

### Plasma MAPK14 quantification by ELISA

Plasma MAPK14 concentrations were measured using a Human MAPK14 ELISA Kit (Finetest; EH2437-96TESTS) at baseline and on postoperative days 2 and 4 in a cohort comprising six patients per group. All RT-qPCR and ELISA assays were performed in technical duplicate. Samples showing substantial variation between duplicate measurements were excluded during quality control, resulting in reduced sample numbers for COX6C and COX7B analyses in the POAF group and for certain ELISA time points. RT-qPCR data were analysed using ΔCq values. Differences between groups were assessed using unpaired two-tailed t-tests with Welch’s correction. To account for multiple comparisons across the four target genes, p-values were adjusted using the two-stage step-up method of Benjamini, Krieger and Yekutieli with a false discovery rate (FDR) of 5% (Q = 0.05).. ELISA data were analysed in R using the lme4, lmerTest, and emmeans packages. MAPK14 concentrations were log-transformed to address positive skewness and analysed using a linear mixed-effects model (LMM) estimated by restricted maximum likelihood (REML). The model included fixed effects for group (POAF vs no-POAF), timepoint (baseline, day 2, and day 4), and their interaction, with subject included as a random effect; timepoint was treated as a categorical factor. Degrees of freedom and P values were calculated using the Kenward-Roger approximation. Model assumptions were assessed by inspection of residual Q-Q plots and histograms overlaid with theoretical normal curves. Post hoc pairwise comparisons between POAF and no-POAF groups at each timepoint were performed using the emmeans package with Šídák adjustment for multiple comparisons. Model estimates and confidence intervals were back-transformed and reported as ratios of geometric means. All statistical analyses were performed in GraphPad Prism (version 11.0.0) and R, with two-tailed P values <0.05 considered statistically significant.

### Peripheral blood cell counts

As part of standard postoperative care, full blood counts were performed using EDTA-preserved blood on Days 1, 2, and 4. Peripheral monocyte, neutrophil, and total white blood cell counts were analysed on a Sysmex XN analyser at a local, UKAS-accredited clinical laboratory (9785) as part of the full blood count. This data was subsequently extracted and used for comparison between the no POAF and POAF groups. Between-group differences at each timepoint, and within-group differences over time, were assessed using a mixed-effects model with Tukey’s multiple comparisons test in GraphPad Prism (version 11.0.0).

## Results

Given the modest cohort size, these findings should be interpreted as exploratory and hypothesis-generating.

nCounter screening identified MAPK14, COX6C and COX7B as higher in patients who developed POAF, and PTPN12 as lower (Fig. 1a). None of these differences remained statistically significant after FDR correction, which is expected given the size of the discovery cohort (n=10 per group), but MAPK14, COX6C and COX7B were carried forward for validation because they linked a differential expression signal to a biologically plausible stress-response and mitochondrial pathway.

**Figure 1:**
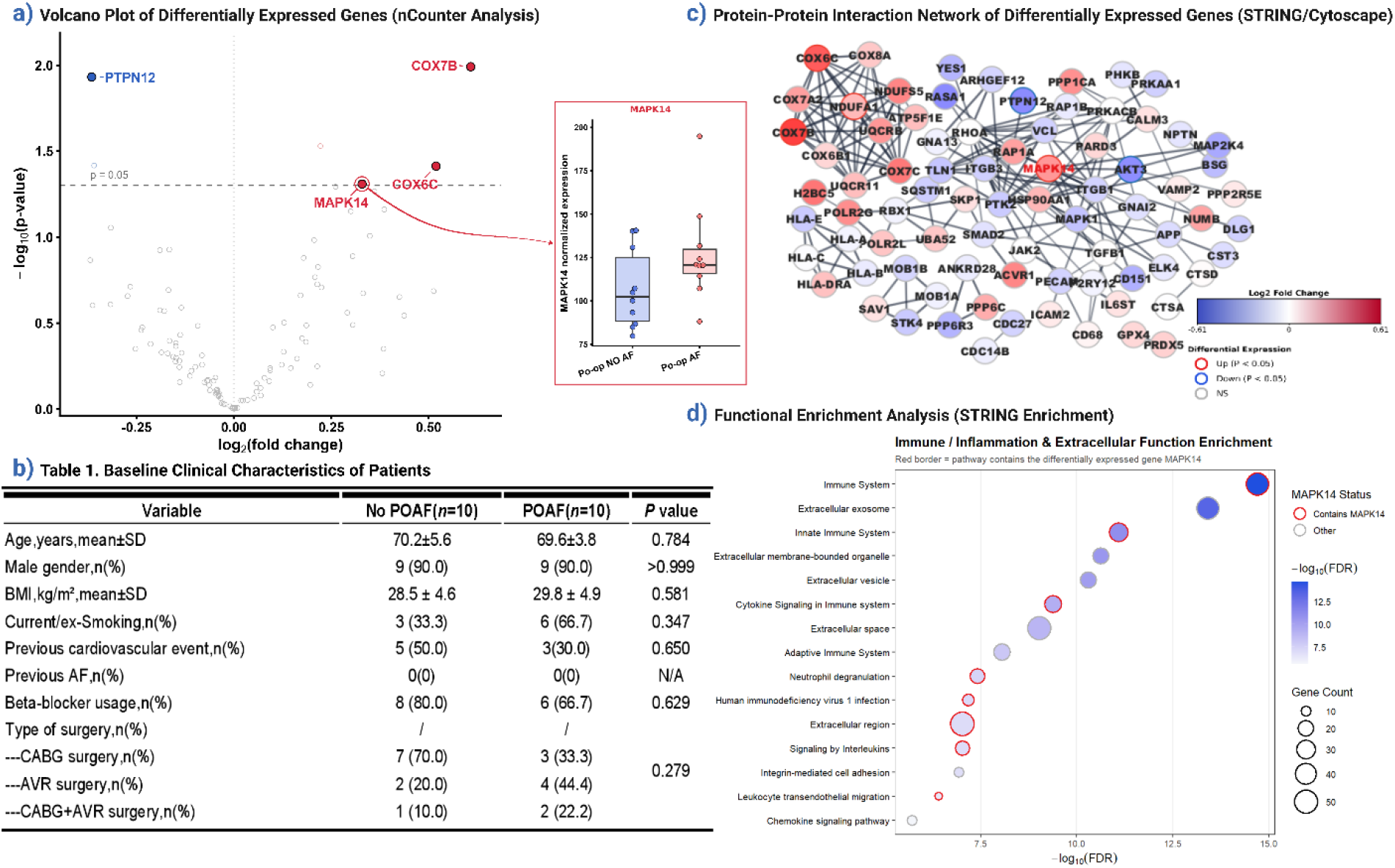
Discovery-stage nCounter transcriptomic analysis highlights MAPK14-related inflammatory signalling as a candidate pathway associated with postoperative atrial fibrillation (Created in BioRender. HOU, JQ. 2026). **(a)**: Plasma nCounter transcriptomic profiling in patients who developed postoperative atrial fibrillation (POAF) and those who remained in sinus rhythm (n=10 per group). The volcano plot displays log2(fold change) versus −log10(P value), with the dashed line indicating nominal P=0.05. Owing to the limited sample size, no transcripts remained significant following false discovery rate (FDR) corrections. Candidate genes selected for downstream investigation based on nominal differential expression and biological relevance are highlighted, including MAPK14, COX6C and COX7B, which showed higher expression in POAF, and PTPN12, which showed lower expression. The inset displays normalised MAPK14 counts, demonstrating a trend towards higher expression in POAF. Genes are shown at nominal significance, and none passed FDR correction. **(b)**: Baseline clinical and operative characteristics of the study groups. Measured variables were broadly comparable between groups (all P values > 0.05). Denominators were 9 for selected variables because of missing clinical records. **(c)**: STRING/Cytoscape protein-protein interaction network generated from nominally differentially expressed genes, illustrating MAPK14 within a connected network of inflammatory, stress-response and signal-transduction pathways. **(d)**: Functional enrichment analysis of nominally differentially expressed genes. Bubble size represents the number of genes contributing to each pathway, colour intensity represents −log10(FDR), and red outliers indicate pathways containing MAPK14.

Baseline clinical characteristics were broadly similar between patients with and without POAF (Fig. 1b). Although the modest sample size limits the ability to exclude subtle baseline differences, the transcriptomic findings do not appear to be explained by major imbalances in the measured variables.

In the STRING/Cytoscape network, these candidates sat within a connected cluster rather than appearing as isolated nodes (Fig. 1c), and functional enrichment of the differentially expressed genes was dominated by immune and inflammatory categories, alongside extracellular process terms (Fig. 1d).

RT-qPCR confirmed the direction of the screening result for three of the four candidates: MAPK14, COX6C and COX7B were all higher in patients with POAF (FDR-corrected q=0.0094, 0.0060 and 0.0094, respectively), while PTPN12 showed a similar trend to nCounter, but was not significantly different (q=0.088) (Fig. 2a). COX6C and COX7B analyses were based on fewer POAF samples (n = 4) than no-POAF samples (n = 6) owing to quality-control exclusions, and this difference in sample size should be considered when interpreting the statistical significance of these findings.

**Figure 2:**
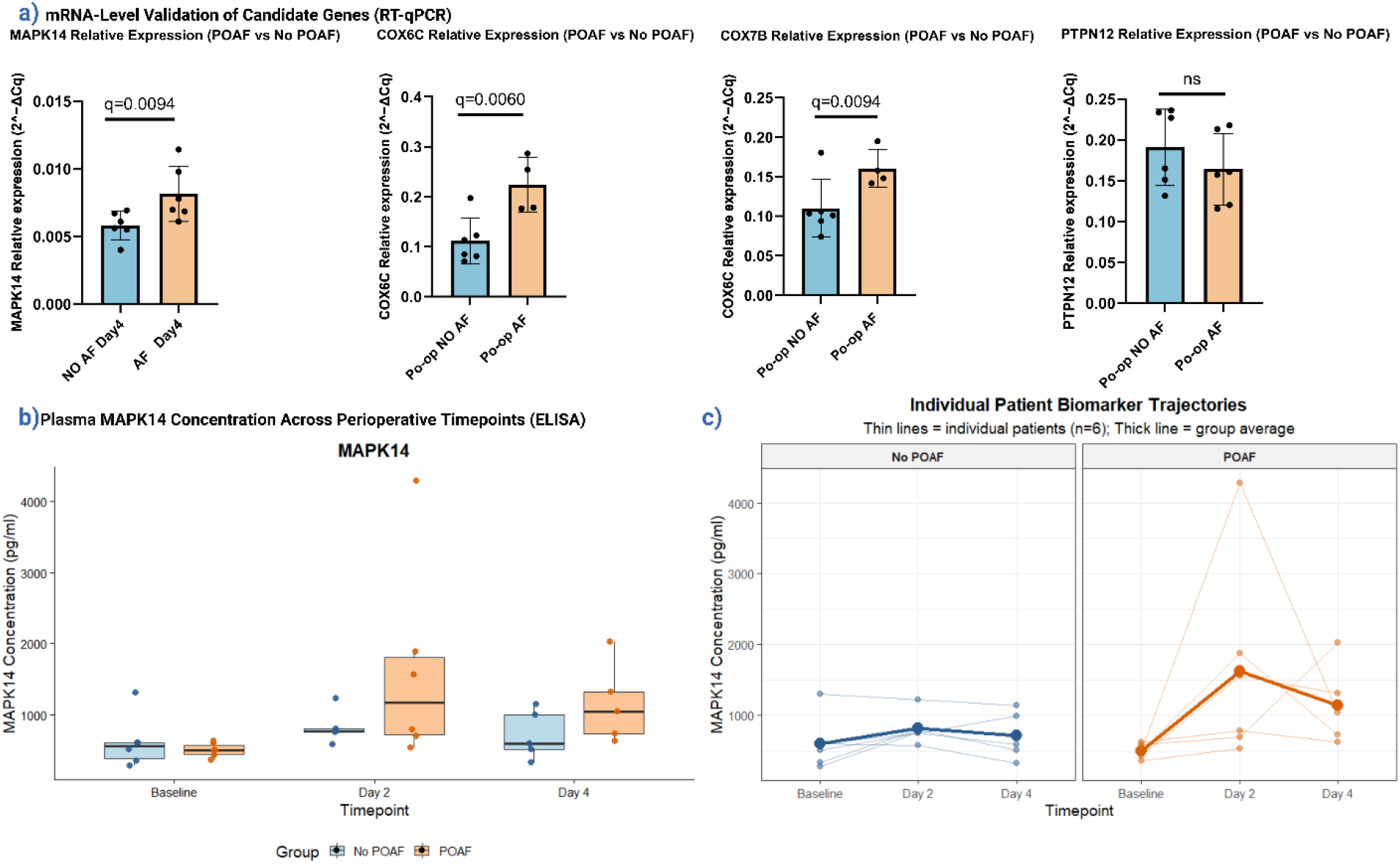
RT-qPCR validation and exploratory longitudinal MAPK14 ELISA readouts in postoperative atrial fibrillation (Created in BioRender. HOU, JQ. 2026). **(a)**: RT-qPCR validation of selected candidate transcripts identified from the exploratory nCounter analysis. Relative expression was calculated using the 2^−ΔΔCt^ method. MAPK14 and PTPN12 were analysed with n=6 per group; COX6C and COX7B were analysed with n=6 in the no-POAF group and n=4 in the POAF group after quality-control exclusions. MAPK14, COX6C and COX7B were higher in POAF (FDR-corrected q=0.0094, 0.0060 and 0.0094, respectively), whereas PTPN12 showed the same expression trend as the nCounter results but was not significantly different. Data are shown as mean±SD with individual points. **(b)** and **(c)**: Exploratory longitudinal plasma MAPK14 ELISA measurements at baseline, postoperative day 2 and postoperative day 4. Plasma MAPK14 concentration is shown for no-POAF and POAF groups. Sample numbers per timepoint were baseline 6/6, day 2 5/6 and day 4 5/5 for no-POAF/POAF, respectively. Individual samples, group means and medians are shown in (c). Given the limited sample size, these analyses should be interpreted as exploratory and hypothesis-generating.

Plasma MAPK14 concentrations increased following surgery in both groups during the first four postoperative days. Although the POAF group showed the greatest increase at postoperative day 2, this difference was not statistically significant, likely reflecting the small sample size at each time point and the reduction in analysable samples following quality-control exclusions.

ELISA measurements were performed in duplicate, and samples showing substantial inter-duplicate variability were excluded according to predefined quality-control criteria. Although plasma MAPK14 concentrations tended to increase more markedly in the POAF group on postoperative day 2, this difference did not remain significant after correction for multiple comparisons.

Peripheral blood monocyte counts were significantly higher in patients who developed POAF than in those who did not, at all three postoperative timepoints (Day 1 P=0.0242, Day 2 P=0.0389, Day 4 P=0.0057; Fig. 3a1). Neutrophil and total white blood cell counts did not differ significantly between the two groups at any timepoint (Fig. 3b1, c1).

**Figure 3:**
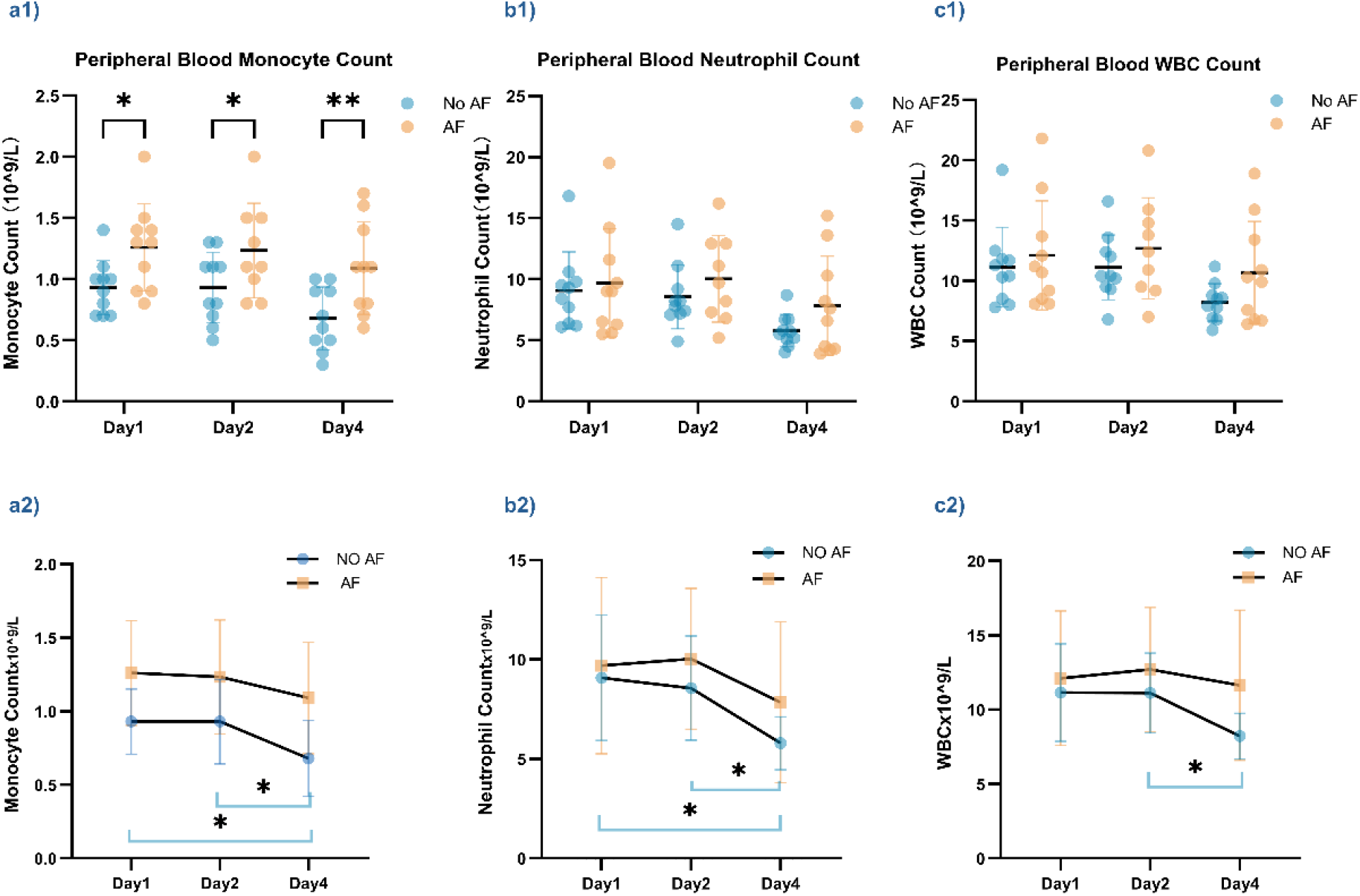
Exploratory analysis of peripheral blood monocyte, neutrophil and white blood cell counts in patients with and without postoperative atrial fibrillation (POAF). **(a1)–(c1):** Peripheral blood monocyte (a1), neutrophil (b1) and total white blood cell (WBC, c1) counts on postoperative day 1, day 2 and day 4, compared between the no-AF and AF groups. Sample numbers per timepoint were day1 10/10, day2 10/9 and day4 10/10 for no-AF/AF, respectively. Data were analysed by mixed-effects model with Tukey’s multiple comparisons test and are shown as individual values with mean±SD. Monocyte counts were higher in the AF group at all three timepoints (day1 P=0.0242, day2 P=0.0389, day4 P=0.0057); neutrophil and WBC counts did not differ significantly between the no-AF and AF groups at any timepoint. **(a2)–(c2):** The same monocyte (a2), neutrophil (b2) and WBC (c2) data re-plotted as group mean±SD across day1, day2 and day4 to illustrate the within-group longitudinal trends. In the no-AF group, monocyte and neutrophil counts declined significantly from day1 and day2 to day4, and WBC counts declined significantly from day2 to day4 (Tukey’s multiple comparisons test following the mixed-effects model); no significant within-group change over time was observed in the AF group for any of the three parameters. *P<0.05, **P<0.01.

When the same data were examined longitudinally within each group, monocyte and neutrophil counts declined significantly from Day 1 and Day 2 to Day 4 in patients without POAF, and white blood cell counts declined significantly from Day 2 to Day 4 in this group; no significant within-group change over time was observed for any of the three parameters in patients with POAF (Fig. 3a2–c2). This sustained elevation of circulating monocytes did not resolve over the first four postoperative days in POAF patients and interestingly provides a cellular correlate to the monocyte/macrophage-related network and pathway signals identified in the transcriptomic screen (Fig. 1c, d).

## Discussion

Targeted transcriptomic profiling using nCounter technology in this small patient cohort identified MAPK14, together with the mitochondrial related transcripts COX6C and COX7B, as candidate biomarkers associated with POAF. These were not isolated signals: in the STRING/Cytoscape network they clustered within a connected inflammatory and stress-response module, and functional enrichment of the differentially expressed genes was dominated by immune and inflammatory categories, consistent with the broader literature linking POAF to inflammatory and fibrotic signalling [5]. This pattern is consistent with previous reports linking POAF to inflammatory and fibrotic signalling [1, 15], and suggests that the screening signal reflects a coordinated response rather than a single gene.

RT-qPCR confirmed higher expression of MAPK14, COX6C and COX7B in patients who developed POAF, and plasma MAPK14 measured by ELISA rose after surgery in both groups, with the largest increase in the POAF group at day 2.

Although the protein-level differences did not remain statistically significant after correction for multiple comparisons, the direction of effect was consistent across the discovery screen, network analysis, RT-qPCR validation and protein measurements. Taken together, the transcriptomic, RT-qPCR and protein-level findings provide convergent support for MAPK14 as a candidate POAF-associated inflammatory signal. These observations are in line with previous evidence implicating macrophage-derived IL-6 signalling in the pathophysiology of POAF [5].

While the molecular analyses implicated MAPK14-associated inflammatory signalling in POAF susceptibility, we next sought evidence that this signal was reflected at the cellular level. To address this, we examined postoperative peripheral blood monocyte, neutrophil and total white blood cell counts.

This cellular evidence was reinforced by routine postoperative blood counts, which demonstrated persistently higher circulating monocyte counts in patients who developed POAF across the first four postoperative days, despite no significant differences in total white blood cell or neutrophil counts between groups. Previous studies have associated higher monocyte counts and monocyte-related inflammatory indices, including the monocyte-to-HDL cholesterol ratio, with an increased risk of POAF and adverse early postoperative outcomes following cardiac surgery [11]. However, most prior investigations have focused on preoperative measurements, and comparatively little is known regarding postoperative monocyte kinetics in relation to POAF occurrence. Our findings therefore extend the existing literature by suggesting that persistence of postoperative monocytosis, rather than a generalised leukocyte response, may characterise the inflammatory milieu associated with POAF. While observational in nature, this interpretation is concordant with the accompanying transcriptomic and protein-level findings implicating MAPK14-associated inflammatory signalling and is consistent with a potential role for monocyte/macrophage-lineage immune pathways in POAF pathophysiology. Nevertheless, causality cannot be inferred, and the relative contribution of other inflammatory cell populations remains uncertain. In addition, postoperative leukocyte profiles are influenced by numerous perioperative factors, including cardiopulmonary bypass exposure, corticosteroid administration, blood transfusion and postoperative complications, all of which may affect circulating monocyte numbers independently of arrhythmia development [16]. Accordingly, these findings should be viewed as hypothesis-generating and require validation in larger, prospectively characterised cohorts incorporating detailed perioperative phenotyping and immune-cell profiling [17].

The nCounter analysis was intended as a discovery screen and identified candidate transcripts associated with POAF; however, the limited cohort size reduced statistical power after correction for multiple testing. Furthermore, RT-qPCR validation was undertaken in the same cohort rather than an independent replication set, and the postoperative increase in MAPK14 measured by ELISA did not remain statistically significant following adjustment for multiple comparisons. Although the present study was not designed to establish a causal role for MAPK14 signalling in POAF, the consistent association observed across our analyses supports its potential relevance to POAF susceptibility. The observed expression patterns are compatible with a role for MAPK14 in the inflammatory response to cardiac surgery and identify this pathway as a promising target for further mechanistic investigation. The next step will be to determine whether MAPK14 and the other candidate markers identified here can be replicated in a larger, independent cohort with sufficient statistical power at each postoperative timepoint. Successful replication would strengthen the evidence linking MAPK14-related signalling to POAF susceptibility and provide a rationale for subsequent studies investigating whether this pathway can be therapeutically modulated. Several anti-inflammatory and autonomic interventions, including colchicine, corticosteroids, pericardial procedures, and autonomic denervation strategies, have already been evaluated in POAF [7–10]. Validation of MAPK14-related signalling could therefore help move beyond observational associations towards the identification of targeted preventive approaches.

## Acknowledgements

RABB acknowledges research funds from the Ellis T Davies Fellowship Endowment and LIV-SRF (Liverpool-Shared Research Facilities) funding from the University of Liverpool.

We thank the LIV-SRF Spatial Profiling Lab for technical support. AD is funded by a J Hill Abram Prize – PhD studentship (award to RABB from the University of Liverpool). G.Y.H.L. has been a consultant and speaker for BMS/Pfizer, Boehringer Ingelheim, Anthos, and Daiichi-Sankyo. No fees are directly received personally. All the disclosures happened outside the submitted work. He is a National Institute for Health and Care Research (NIHR) Senior Investigator Emeritus and co-PI of the AFFIRMO project on multimorbidity in AF (grant agreement No 899871), TARGET project on digital twins for personalized management of atrial fibrillation and stroke (grant agreement No 101136244), and ARISTOTELES project on artificial intelligence for management of chronic long-term conditions (grant agreement No 101080189), which are all funded by the EU’s Horizon Europe Research and Innovation program.

## Author Contributions

RABB conceived the research idea to pursue nCounter plasma studies. GYL and NT designed the AF Big Picture cohort study. BK is the surgeon involved in this study and EH contributed towards sample collection. JH performed experiments and analysed the first data reports. FG, JH, AD, PS, NH, RABB and IMD contributed to RT-qPCR experiments. JH, NT, HP, AD, YM contributed to ELISA experiments. NT, JH, NH and RABB contributed to statistical analysis. RABB and JH conducted literature search for the study and wrote the first manuscript draft. All authors have contributed to the content and refinement of the manuscript. All authors contributed intellectually to the study.

## Figure Legends

**Supplementary Table1.**
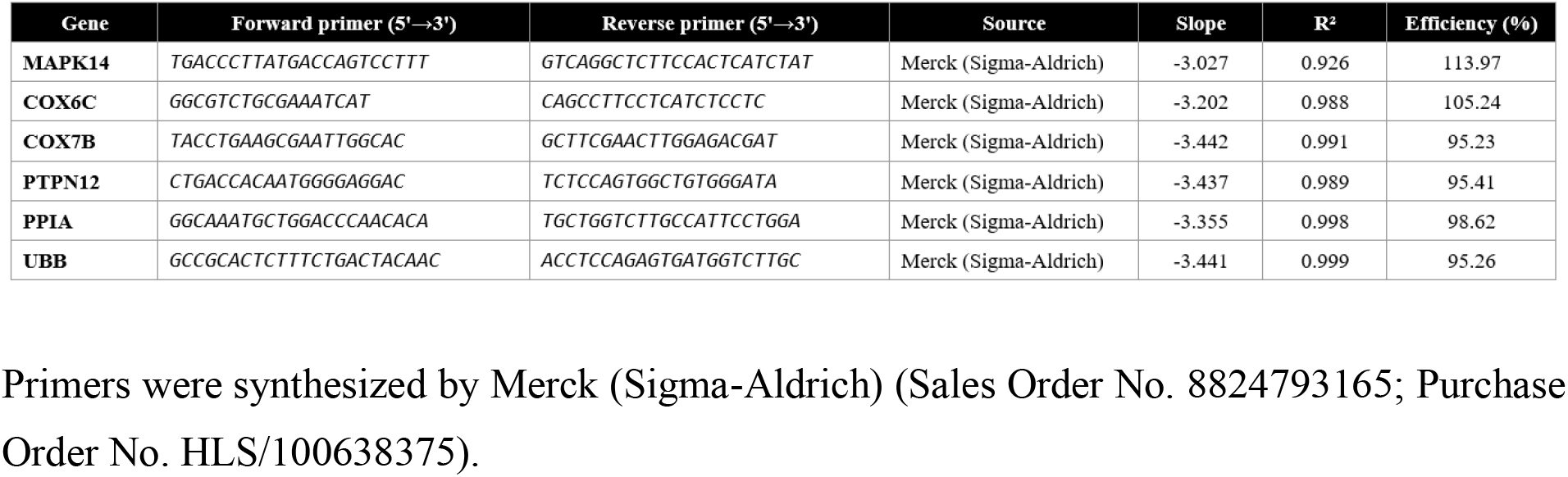
Primer sequences and qPCR validation parameters.

