## Supplemental Table 1 for "Integrated Transcriptomic Profiling Reveals MAPK14-Related Inflammatory Signalling Associated with Postoperative Atrial Fibrillation"

### **Supplementary Table1. Primer sequences and qPCR validation parameters**

| **Gene** | **Forward primer (5'→3')** | **Reverse primer (5'→3')** | **Source** | **Slope** | **R²** | **Efficiency (%)** |
| --- | --- | --- | --- | --- | --- | --- |
| **MAPK14** | *TGACCCTTATGACCAGTCCTTT* | *GTCAGGCTCTTCCACTCATCTAT* | Merck (Sigma-Aldrich) | -3.027 | 0.926 | 113.97 |
| **COX6C** | *GGCGTCTGCGAAATCAT* | *CAGCCTTCCTCATCTCCTC* | Merck (Sigma-Aldrich) | -3.202 | 0.988 | 105.24 |
| **COX7B** | *TACCTGAAGCGAATTGGCAC* | *GCTTCGAACTTGGAGACGAT* | Merck (Sigma-Aldrich) | -3.442 | 0.991 | 95.23 |
| **PTPN12** | *CTGACCACAATGGGGAGGAC* | *TCTCCAGTGGCTGTGGGATA* | Merck (Sigma-Aldrich) | -3.437 | 0.989 | 95.41 |
| **PPIA** | *GGCAAATGCTGGACCCAACACA* | *TGCTGGTCTTGCCATTCCTGGA* | Merck (Sigma-Aldrich) | -3.355 | 0.998 | 98.62 |
| **UBB** | *GCCGCACTCTTTCTGACTACAAC* | *ACCTCCAGAGTGATGGTCTTGC* | Merck (Sigma-Aldrich) | -3.441 | 0.999 | 95.26 |

*Primers were synthesized by Merck (Sigma-Aldrich) (Sales Order No. 8824793165; Purchase Order No. HLS/100638375).*
